# Uncovering methylation-dependent genetic effects on regulatory element function in diverse genomes

**DOI:** 10.1101/2024.08.23.609412

**Authors:** Rachel M. Petersen, Christopher M. Vockley, Amanda J. Lea

**Affiliations:** Department of Biological Sciences, Vanderbilt University, Nashville, TN; Evolutionary Studies Initiative, Vanderbilt University, Nashville, TN; The Broad Institute of MIT and Harvard, Cambridge, MA; Vanderbilt Genetics Institute, Vanderbilt University, Nashville, TN

**Keywords:** mSTARR-seq, massively parallel reporter assay, DNA methylation, allele specific expression, methylation-dependent transcription factor binding

## Abstract

A major goal in evolutionary biology and biomedicine is to understand the complex interactions between genetic variants, the epigenome, and gene expression. However, the causal relationships between these factors remain poorly understood. mSTARR-seq, a methylation-sensitive massively parallel reporter assay, is capable of identifying methylation-dependent regulatory activity at many thousands of genomic regions simultaneously, and allows for the testing of causal relationships between DNA methylation and gene expression on a region-by-region basis. Here, we developed a multiplexed mSTARR-seq protocol to assay naturally occurring human genetic variation from 25 individuals sampled from 10 localities in Europe and Africa. We identified 6,957 regulatory elements in either the unmethylated or methylated state, and this set was enriched for enhancer and promoter annotations, as expected. The expression of 58% of these regulatory elements was modulated by methylation, which was generally associated with decreased RNA expression. Within our set of regulatory elements, we used allele-specific expression analyses to identify 8,020 sites with genetic effects on gene regulation; further, we found that 42.3% of these genetic effects varied between methylated and unmethylated states. Sites exhibiting methylation-dependent genetic effects were enriched for GWAS and EWAS annotations, implicating them in human disease. Compared to datasets that assay DNA from a single European individual, our multiplexed assay uncovers dramatically more genetic effects and methylation-dependent genetic effects, highlighting the importance of including diverse individuals in assays which aim to understand gene regulatory processes.

## INTRODUCTION

A major goal in evolutionary biology, human genomics, and biomedicine is to understand how the genome generates complex traits, which at the molecular level often involves complex interactions between genetic variants, the epigenome, and gene expression. DNA methylation (DNAm) is an epigenetic process whereby a methyl group is added to a CpG dinucleotide motif (this is the most common site of DNAm in mammals). DNAm plays important roles in processes that generate cellular and organismal phenotypic variation by controlling gene expression. For example, the establishment and erasure of DNAm is associated with the development and differentiation of tissues (Luo et al. 2018; Greenberg and Bourc’his 2019; Loyfer et al. 2023), responses to environmental exposures (Waterland and Jirtle 2003; Weaver et al. 2004; Garg et al. 2018; Perera et al. 2020; Anderson et al. 2024), and aging-related disease progression (Jin and Liu 2018; Salameh et al. 2020; Yim et al. 2020).

While DNAm is canonically associated with repressed transcription of nearby genes—for example through alterations in transcription factor (TF) recruitment and/or binding affinity (Holliday and Pugh 1975; Riggs 1975; Han et al. 2011; Yin et al. 2017; Kaluscha et al. 2022)—recent studies have shown that this relationship varies across the genome. For example, in many regions, DNAm is functionally negligible, having little to no effect on TF binding or downstream mRNA production. This has been observed via protein binding microarrays and high-throughput systematic evolution of ligands by exponential enrichment (SELEX) methods that have quantified comparable TF binding preferences to methylated versus unmethylated sequences in vitro (Hu et al. 2013; Kribelbauer et al. 2020). Similarly, studies of primary cells that have employed time course designs or single molecule footprinting have also concluded that much of the genomic variation in DNAm does not directly impact gene expression (Pacis et al. 2015; Kreibich et al. 2023). In addition to many regions being functionally insensitive to methylation, others display DNAm-expression relationships that are opposite the canonical pattern. For example, the positioning of methylation surrounding TF binding motifs can result in increased binding affinity or altered chromatin conformation which can ultimately result in higher levels of DNAm correlating with increased expression of nearby genes (Luo et al. 2021; Monteagudo-Sánchez et al. 2024). Given the complexities of DNAm-expression relationships, it is therefore essential to directly evaluate whether and how DNAm impacts local gene transcription, rather than assuming a functional (typically repressive) relationship.

In addition to DNAm, it is also clear that genetic variation impacts cis gene expression, often through similar processes such as variable TF recruitment and binding (Kasowski et al. 2010; Deplancke et al. 2016). Accordingly, expression quantitative trait loci (eQTL) studies have documented a myriad of associations between cis genetic variants and gene expression in human populations (GTEx Consortium 2020; Kim-Hellmuth et al. 2020; Kerimov et al. 2021; Zeng et al. 2022), with high powered studies now reporting ≥1 cis eQTL for essentially every gene in the genome (Gong et al. 2018; Võsa et al. 2021). Further, recent work suggests that genotype-expression relationships may in some cases depend on the local methylation context: Zeng and colleagues mapped 1,731 “methylation-dependent eQTL” (where the eQTL presence depends on local methylation levels): the vast majority of these eQTL had not previously been reported as standard eQTL. Mechanistically, the authors hypothesized that methylation may induce alterations to TF recruitment and 3D chromatin structure that modulates genotype-expression relationships (Zeng et al. 2023). Although studies such as these provide correlational evidence that DNAm may alter genotype-expression relationships in important ways (Banovich et al. 2014; Singh et al. 2020), experimental studies demonstrating cause and effect relationships are lacking.

To address these gaps, massively parallel reporter assays (MPRAs) have become a useful tool for experimentally testing the causal effects of cis genetic variation and DNAm on transcriptional activity. This family of assays allows the user to test millions of fragments for regulatory activity, in the presence of genetic variation or experimentally induced DNAm, in a single experiment (Inoue and Ahituv 2015; Vockley et al. 2015; Tewhey et al. 2016; Ulirsch et al. 2016; Lea et al. 2018). The original MPRA design involved cloning a potential regulatory region of interest into a plasmid backbone upstream of a reporter gene and a unique sequence-associated barcode (Patwardhan et al. 2009; Kwasnieski et al. 2012; Melnikov et al. 2012; Patwardhan et al. 2012; Sharon et al. 2012). Plasmids are transfected into a cell line, incubated, and regulatory activity of the insert is then quantified via barcode sequencing. These methods have been used to test hundreds of thousands of regions of the genome for activity, as well as the effects of polymorphisms within regulatory regions on activity (for example: Tewhey et al. 2016; Kalita et al. 2018; Griesemer et al. 2021; Jagoda et al. 2022; Siraj et al. 2023).

Building upon these methods, self-transcribing active regulatory region sequencing (STARR-seq) was developed as a type of MPRA; in this assay, regions of interest are inserted within the 3’ UTR, such that active regulatory elements drive expression of transcripts that include the focal sequence itself. Regulatory activity then directly scales with plasmid-derived mRNA levels of the focal sequence (Arnold et al. 2013). Finally, methyl-STARR-seq (mSTARR-seq) pairs the STARR-seq approach with methylation manipulation of query fragments to test for the effects of DNAm on regulatory activity (Lea et al. 2018). Consistent with other techniques, mSTARR-seq studies have shown that experimental manipulation of DNAm does not always impact gene expression in the canonical direction: ∼½ to ¾ of tested regulatory elements were found to be insensitive to changes in DNAm, and 2-14% of regions with methylation-dependent (MD) regulatory function actually exhibit *increased* transcriptional activity following experimental methylation (Lea et al. 2018; Johnston et al. 2024). Consistent with previous studies, MD regulatory elements were enriched for binding motifs of TFs known to preferentially bind to methylated DNA (Hu et al. 2013; Zuo et al. 2017; Luo et al. 2021).

While MPRA methods excel in directly testing for causal relationships between genetic or epigenetic variation and regulatory activity, studies of genetic variation have almost exclusively focused on testing synthesized sequences (Tewhey et al. 2016; Ulirsch et al. 2016; Kalita et al. 2018; Siraj et al. 2023), which are expensive to produce, commonly short (∼180bp) in length, and may lack naturally occurring patterns of linkage disequilibrium (LD); in turn, studies of epigenetic variation have exclusively focused on DNA from a single European individual (Lea et al. 2018; Johnston et al. 2024). The common practice of benchmarking experimental assays on a single European genome limits our knowledge of how genetic variation impacts functional genomic mechanisms and perpetuates the underrepresentation of non-European individuals in genomic research (Gurdasani et al. 2019; Mills and Rahal 2020). To improve upon these shortcomings, we developed multiplexed mSTARR-seq, an extension of mSTARR-seq that can incorporate genetically diverse input and is thus capable of experimentally testing for interactions between DNAm and genotype in high throughput.

Applying this method, we created a genetically heterogeneous, barcoded DNA input library from 25 individuals from the 1000 Genomes Project originating from 10 different populations throughout Europe and Africa (The 1000 Genomes Project Consortium 2015). This multiplexed library covered 36.1 M unique DNA fragments, 1.95 Gb of genomic sequence, 19.6M unique genetic variants, and 23.2 M unique CpG sites. Genetic variation within our sample allowed us to perform allele specific expression (ASE) analyses to identify allelic biases in regulatory function in both methylated and unmethylated contexts. In this manner, we experimentally determined how methylation and genetic variation can interact to modify regulatory activity on a locus-by-locus basis at hundreds of thousands of sites throughout the genome simultaneously. Using these methods, we identify hundreds of methylation-dependent genetic effect sites, and explore where they are located throughout the genome, their proximity to TF binding motifs, and their overlap with genome-wide association study (GWAS) and epigenome-wide association study (EWAS) hits. Lastly, we demonstrate how the multiplexing of DNA from genetically diverse individuals vastly increases mSTARR-seq’s ability to identify genetic effects as well as interactions between genetic and epigenetic effects, a benefit that is magnified especially by the inclusion of individuals of African origin. As the field of genomics works toward characterizing human genetic diversity and its impacts on regulatory function, multiplexed assays such as this will be an important tool to maximize the breadth of sequence variation that can be investigated and contribute to a more comprehensive understanding of genome regulation.

## RESULTS

### Multiplexed mSTARR-seq produces a genetically diverse input library

We extracted DNA from lymphoblastoid cell lines (LCLs) of 25 different 1000 Genomes individuals originating from 10 different populations (The 1000 Genomes Project Consortium 2015; Figure 1A; Table S1) and digested each individual’s DNA using the Msp1 restriction enzyme, whose recognition site is CCGG, to enrich for CpG sites. Similar to previous mSTARR-seq experiments (Lea et al. 2018; Johnston et al. 2024), we performed PCR on NEB DNA libraries using a primer which joins to the adapter and creates a 15 bp overhang complementary to the mSTARR plasmid for Gibson assembly. New to this protocol, the R primer binding to the 3’ end of the fragment also contained a P7 sequence and a CpG free 6 bp barcode, situated in between the sequences complementary to the NEB adapter and the plasmid backbone (Table S2; Figure S1A). After confirming our design (see SI text for methods, Figure S2), we pooled barcoded DNA libraries from different individuals to create two diverse input pools, with many of the same individuals (n=16) present in both pools but harboring differing barcodes (Figure S1B; Table S3). By sequencing most individuals twice with different barcodes, our goal was to limit any potential impacts that the barcode sequence (which resides in the query fragment) may have on our results. We cloned our genetically diverse input libraries into the mSTARR vector, performed multiple large scale bacterial transformations, and confirmed the diversity of the resulting pool (Figure S1C & S3). We proceeded following Lea et al. 2018 to enzymatically methylate half of the plasmids and expose the other half to a mock/sham control (Figure S1D). We then transfected methylated and unmethylated plasmids into 6 replicate flasks each using the K562 myeloid cell line (Figure S1E). Following a 48-hour incubation, we co-extracted DNA and RNA, sequenced the resulting replicate-level DNA and RNA libraries (Table S4), and assigned reads to individuals within each replicate through the demultiplexing of individual-specific barcodes (Figure 1B).

**Figure 1.**
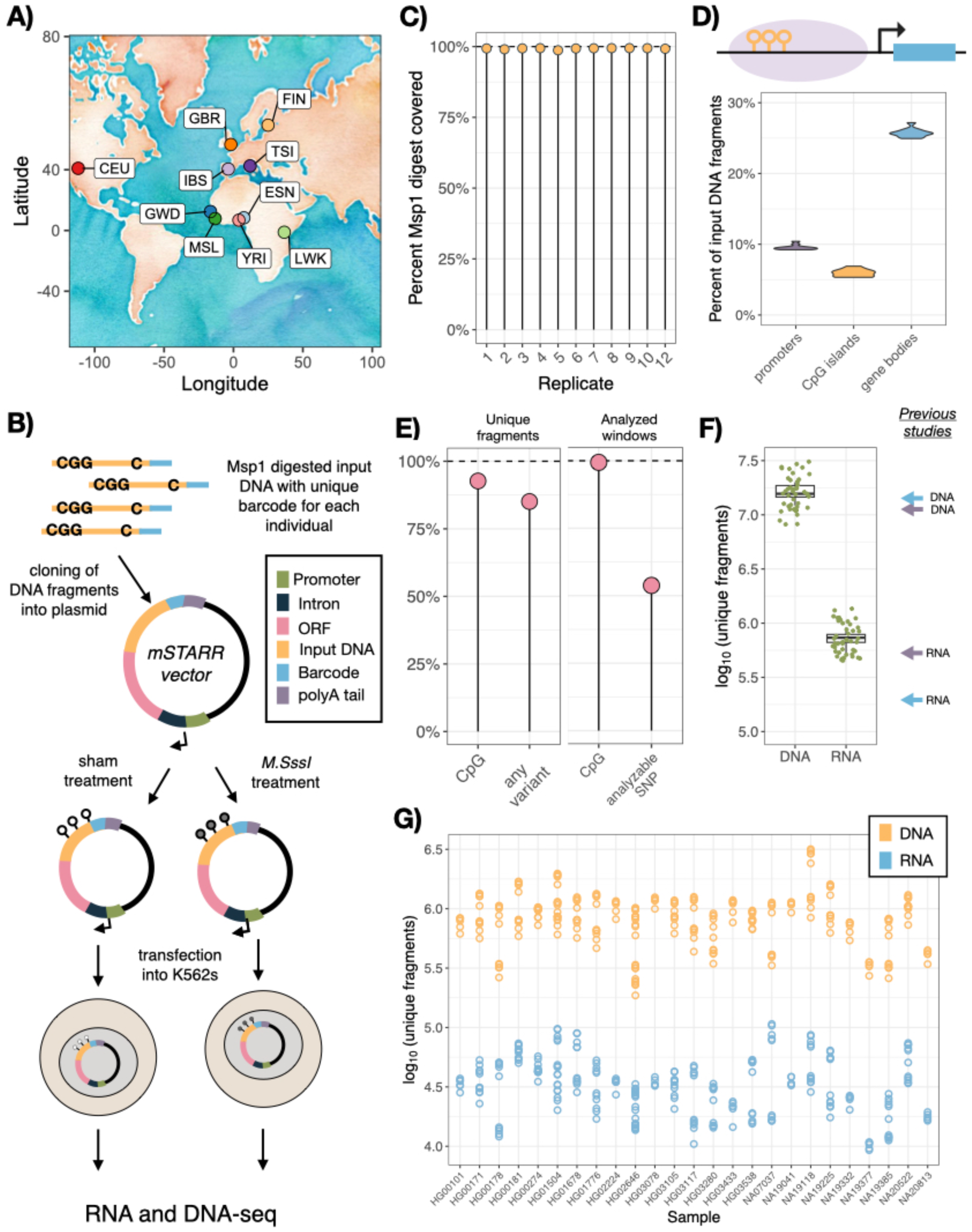
Multiplexed mSTARR-seq assays a diverse input library. A) Sampling locations of the 25 individuals included in the assay: CEU= Utah residents (CEPH), ESN= Esan in Nigeria, FIN= Finnish in Finland, GBR= British in England and Scotland, GWD= Gambian in Western Divisions in the Gambia, IBS= Iberian population in Spain, LWK= Luhya in Webuye, Kenya, MSL= Mende in Sierra Leone, TSI= Toscani in Italy, YRI= Yoruba in Ibadan, Nigeria. B) multiplexed mSTARR-seq design: sample-specific barcodes are added to Msp1 digested input DNA and inserted into the mSTARR vector downstream of a promoter, intron, and open reading frame (ORF). Plasmids are exposed to a methylation treatment or sham control and transfected into K562 cells, incubated for 48 hrs, and DNA and RNA is extracted and sequenced. C) percentage of an in silico Msp1 digest of the human genome that is represented in the DNA input of each replicate, D) percentage of input DNA fragments located within promoters, CpG islands, and gene bodies in each replicate (note these are not mutually exclusive annotations), E) Percent of unique DNA fragments that contain at least one CpG site and at least one SNP (left panel), and the percent of analyzed windows (n=525,074) that contain at least one CpG site and at least one analyzable SNP (i.e., biallelic, > 0.05 MAF, and was called in our joint genotyping analysis; right panel), F) number of unique DNA and RNA fragments observed in each replicate; the mean number of fragments included in each replicate in Lea et al. 2018 (purple arrows) and Johnston et al. 2024 (blue arrows) are shown for comparison, G) number of unique DNA and RNA fragments included in each replicate from each of the 25 individuals included in the assay.

The DNA input fragments covered the majority of an in silico Msp1 digest of the human genome: each replicate covered >98% of the expected fragments (Figure 1C) and each individual within each replicate covered on average 91.3% of the expected fragments. 84.9% of reads started or ended with an Msp1 cut site. As expected, DNA input fragments were enriched for functionally important areas of the genome, with 41% located within protein coding sequences, 18% in promoter regions, and 17% in CpG islands (Figure 1D). In total, the assay included 36.1 M unique fragments, covering 1.95 Gb of unique sequence; these fragments included 23.2 M unique CpG sites and 19.6M unique genetic variants, 737,394 of which were biallelic SNPs with a minor allele frequency (MAF) > 0.05 that could be called within our dataset via joint genotyping. 92.7% of fragments contained at least 1 CpG site and 85.1% of fragments contained at least 1 SNP (Figure 1E). After restricting to genomic windows with sufficient coverage across replicates to facilitate statistical modeling and to SNPs called within our libraries (described below and in Methods), 99.4% of windows contained at least 1 CpG site, totaling 6.2 M CpGs sites, and 53.8% of windows contained at least 1 analyzable SNP (i.e., biallelic, > 0.05 MAF, and called in our joint genotyping analysis), totaling 282,403 SNPs (Figure 1E). Detailed information on sequencing depth can be found in Table S4.

As observed in previous STARR-seq and mSTARR-seq work (Arnold et al. 2013; Vockley et al. 2015; Lea et al. 2018; Johnston et al. 2024), many input DNA fragments do not possess regulatory activity and thus do not generate plasmid RNA output. We found that each replicate contained 736,006 ± 32,831 unique RNA fragments (generated from 15.9 ± 0.95 M unique DNA fragments). Importantly, the average complexity of both our RNA and DNA libraries slightly exceeds that observed in previous mSTARR experiments (Figure 1F; Lea et al. 2018; Johnston et al. 2024). All individuals achieved comparable representation across replicates, with a mean of 35,047 ± 2,707 unique RNA fragments generated from 896,899 ± 58,144 unique DNA fragments per individual per replicate (Figure 1G). The number of windows/variants tested at each data analysis step can be found in Figure S4.

### Multiplexed mSTARR-seq identifies methylation-dependent regulatory elements

To identify generalizable regulatory and MD regulatory elements, we combined uniquely mapped reads from all individuals within a replicate, overlapped these with 400 bp non-overlapping genomic windows, and filtered for windows with adequate coverage (≥ 1 DNA read in half of replicates from both conditions, ≥ 1 RNA read in half of replicates in either condition, as in Lea et al. 2018 and Johnston et al. 2024). Following filtering for DNA and RNA coverage, we were left with 525,074 400 bp windows, of which 522,294 (99.4%) contained at least 1 CpG site. We tested for regulatory activity at each 400bp window in methylated and unmethylated conditions separately by asking whether the abundance of the reporter-gene derived mRNA exceeded the amount of input DNA for that window (Figure 2A). We identified 6,221 windows with regulatory activity in the unmethylated replicates and 2,513 windows with regulatory activity in the methylated replicates (at a 1% FDR), representing 1.18% and 0.48% of tested fragments in the unmethylated and methylated condition respectively (Figure 2B). When we processed data from the current study and two previous mSTARR-seq studies using identical pipelines (Lea et al. 2018, Johnston et al. 2024), we found a similar proportion of tested windows displaying regulatory activity (unmethylated condition: 0.78-1.9%, methylated condition: 0.14-1.7%; Table S5), a significant correlation in effect sizes within regulatory windows (Pearson’s correlation: R=0.48-0.60, Figure S5 & S6), and a strong enrichment for shared identification of regulatory windows across both conditions (Fisher’s exact test: OR= 23.3-79.65; Table S5). As expected, the regulatory windows we identified were significantly enriched for strong enhancer and active promoter chromHMM annotations in K562s (Figure 2C; Table S6). When comparing regulatory windows that were found in one condition but not the other, we found that regulatory windows specific to the unmethylated condition were particularly enriched for promoter and enhancer annotations, while those that were specific to the methylated condition were more enriched for heterochromatin and regions of weak transcription (Figure S7; Table S7)

**Figure 2.**
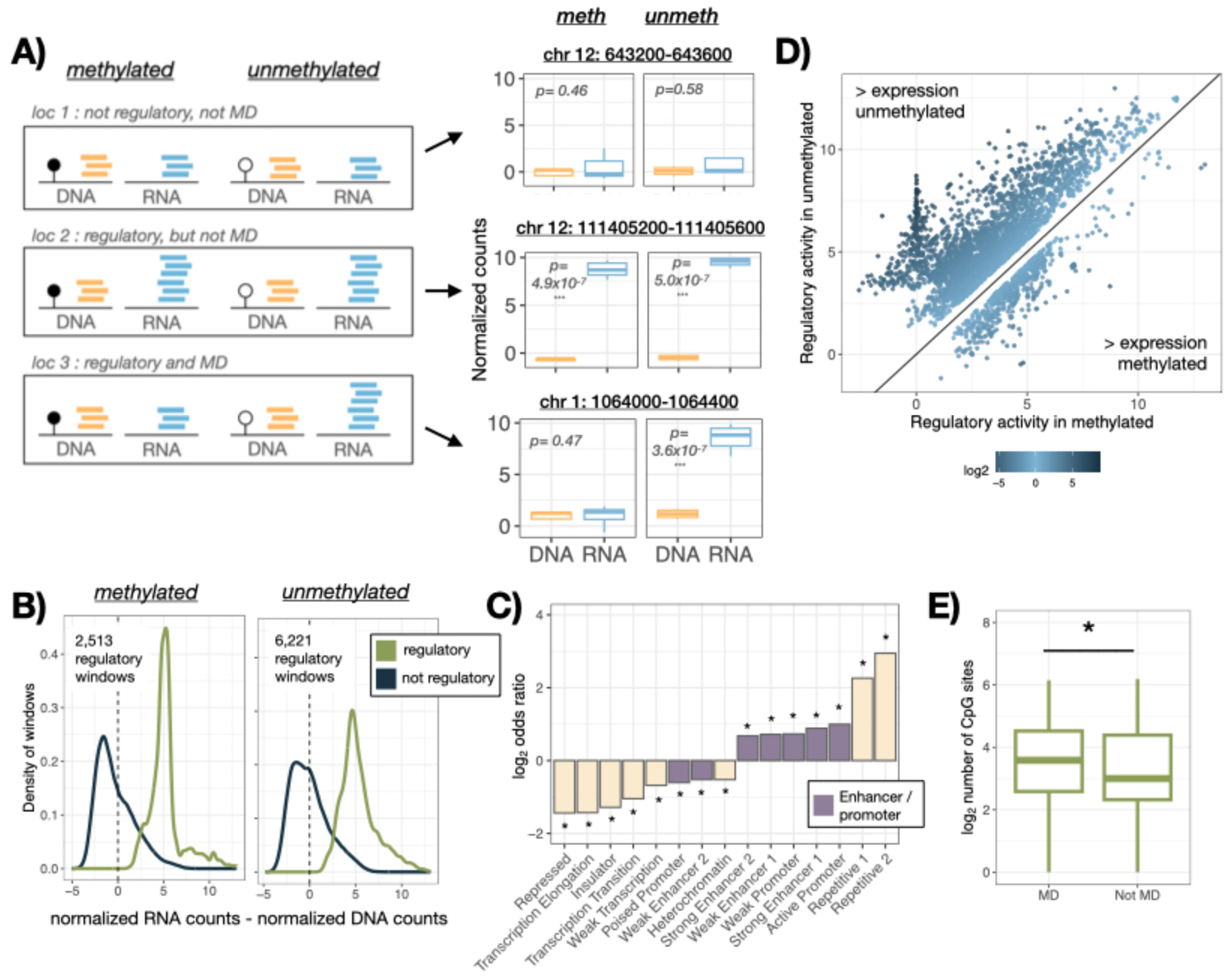
Multiplexed mSTARR-seq identifies regulatory and methylation-dependent regulatory activity. A) Heuristic patterns of read pileups associated with the identification of regulatory activity and methylation-dependence, with a example of the normalized read counts for a window falling into each of these categories, B) density of regulatory and non-regulatory windows (area under each curve normalized to 1) in relation to the difference between normalized RNA and DNA counts for that window C) fisher’s exact test for enrichment in chromHMM genomic annotations when comparing windows with regulatory activity (combined across conditions) versus non-regulatory windows, bars above y=0 indicate annotations that are over enriched in regulatory windows and bars below y=0 indicate annotations that are under enriched in regulatory windows, purple bars indicate enhancer and promoter annotations and stars indicate significant over/under enrichment of that annotation type (full results are presented in Table S6), D) regulatory activity in the methylated versus unmethylated condition for MD windows colored by the log FC between conditions: 84.8% of MD windows have greater activity in the unmethylated condition, with the clustering of sites at x=0 representing windows whose expression is entirely repressed by methylation, E) windows with MD regulatory activity have a greater number of CpG sites than regulatory windows that are not modulated by methylation.

To test for methylation-dependence in regulatory activity, we used a Multivariate Adaptive Shrinkage approach (mashr) (Urbut et al. 2019), which enables testing for effect size sharing versus heterogeneity across conditions. We defined MD regulatory elements as those exhibiting regulatory activity in at least 1 condition (local false sign rate: LFSR <0.05) and with a log fold change (logFC) in regulatory activity > 1.5 between conditions (Figure 2A). Of the 6,957 unique windows with regulatory activity in one condition, 4,052 displayed methylation-dependent activity, meaning that methylation modulated the degree to which the fragment drove gene expression for 58.2% of tested fragments. After rerunning previous mSTARR-seq datasets using this pipeline, we find a similar proportion of regulatory windows exhibiting methylation-dependence (Johnston et al. 2024: 67.6%, Lea et al. 2018: 80.2%, Table S5). As expected, the majority of windows (84.8%) had greater transcriptional activity in the unmethylated relative to the methylated condition (Figure 2D). As noted in previous work, we found that windows displaying MD regulatory activity had a greater number of CpG sites relative to regulatory elements that were not modulated by methylation (T-test: t= 6.38, df= 6352.2, p= 1.9×10^−10^; Figure 2E). Further, methylation-dependent windows were very unlikely to be CpG free relative to the background set of all regulatory windows tested (Fisher’s test: OR= 0.38, p= 5.76×10^−6^).

### Genetic variation impacts regulatory function

To interrogate the effect of genotype on regulatory activity, we estimated allele specific RNA output versus DNA input in the methylated and unmethylated conditions separately, using standard pipelines in GATK (Van der Auwera er al. 2013). Our entire dataset included 737,394 biallelic variants with a MAF > 0.05. After filtering for coverage and for windows with only 1 SNP or haplotype that were in active regulatory windows, we retained 8,570 and 5,853 variants for the unmethylated and methylated conditions for downstream analyses. 3,636 and 2,816 of these variants were monoallelically expressed in a given condition (923 shared between conditions), and thus exhibited extremely strong ASE (see Methods). The number of sites retained following each filtering step are presented in Table S8. For sites that were not monoallelically expressed, we used beta binomial modeling (Lesnoff and Lancelot 2012) to test whether the proportion of reference to alternate allele counts significantly differed between the DNA input and the mRNA output at each site (Figure 3A). With this method, we discovered 2,807 sites with ASE at a 1% FDR threshold: 1,858 sites in the unmethylated state (37.6% of tested sites), and 1,150 sites in the methylated state (37.9% of tested sites; Figure 3B). Sequencing depth did not significantly differ between ASE and non-ASE sites, suggesting that the identification of ASE is not due to variability or noise associated with lower DNA read counts (T-test: unmethylated replicates: t= −0.278, df= 4112.2, p=0.78; methylated replicates: t= 1.18, df= 2732.4, p=0.24; Figure S8).

**Figure 3.**
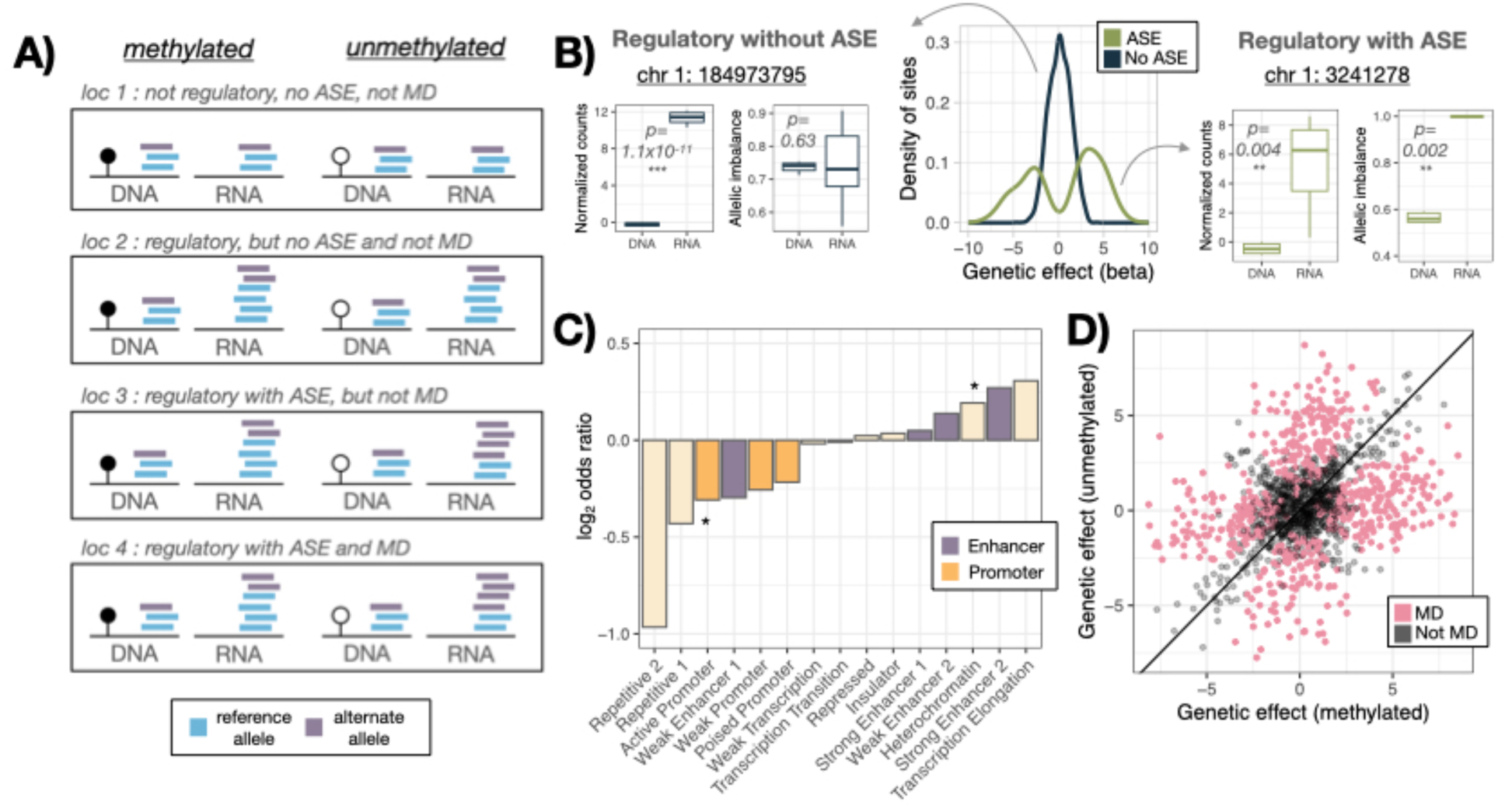
Multiplexed mSTARR-seq identifies allele specific regulatory activity that is modulated by methylation. A) Patterns of read pileups associated with the identification of regulatory activity, ASE, and MD ASE, reference alleles indicated as dark colors, alternate alleles indicated as light colors, B) density of tested sites with ASE versus without ASE in the unmethylated condition, with an example plot of each showing regulatory activity (normalized DNA and RNA counts) and ASE (the ratio of reference allele to total counts, i.e., allelic imbalance) present in each replicate, C) fisher’s exact test for enrichment in chromHMM genomic annotations when comparing ASE sites (combined across conditions) versus non-ASE sites, bars above y=0 indicate annotations that are over enriched in ASE sites and bars below y=0 indicate annotations that are under enriched in ASE sites, purple bars indicate enhancer annotations, orange bars indicate promoter annotations, and stars indicate significant (p < 0.05) over/under enrichment of that annotation type (full results are presented in Table S9), D) the genetic effect in the methylated condition plotted against the genetic effect in unmethylated condition for ASE sites, with the 575 MD ASE sites highlighted in pink.

Recent work has highlighted that phenotypically relevant variants identified via GWAS are typically located in non-coding regions distal to TSSs and promoter regions (Maurano et al. 2012; Mostafavi et al. 2023). Conversely, eQTL signals commonly cluster within promoter regions near TSSs, and may be less relevant to complex traits as they often fail to colocalize with GWAS hits (Chun et al. 2017; GTEx Consortium 2020; Mostafavi et al. 2023). Thus, we aimed to explore where in the genome variants exerting an allele specific effect on regulatory activity were located by comparing these to SNPs that were located within regulatory regions but did not exert allele specific effects on expression. To do so, we combined sites with significant ASE (in our beta binomial model) with those exhibiting monoallelic expression (a particularly strong form of ASE) in either the methylated or unmethylated condition; we compared their proximity to the nearest TSS and their genomic annotations with regulatory variants that did not display allele specific effects. Similar to GWAS sites, we found that variant sites displaying ASE were located farther from the nearest TSS (T-test: t= −4.19, df= 8284.8, p=2.83×10^−5^, Figure S9), and were less likely to be located within active promoter regions (Fisher’s test: OR=0.81, p=0.0004; Figure 3C). Instead, sites with allele specific regulatory effects were more likely to be located in enhancer annotations (although this trend does not reach statistical significance; Figure 3C; Table S9). These results are robust to the exclusion of monoallelic sites in our test dataset (Figure S10, see SI text for full results).

### Methylation impacts allele specific regulatory function

To understand how methylation directly impacts ASE, we again used mashr to compare effect size estimates from our beta binomial model between conditions, and asked whether the genetic effect on regulatory activity is modified by experimental methylation. We considered a SNP to have MD genetic effects if it exhibited ASE in at least 1 condition (LFSR <0.05) and the logFC between conditions for the genetic effect was > 1.5. We tested 1,359 sites with ASE for methylation-dependence, and found 575 sites with MD genetic effects (43.2% of tested sites, Figure 3E). 208 of the tested sites directly overlapped with either the C or G of a CpG site in the hg38 reference genome, such that methylation could only occur for one genotype. Of these 208 CpG disrupting sites, 74% exhibited MD genetic effects (n= 154 sites).

For SNPs that did not disrupt a CpG site (n= 421), we explored which condition was associated with greater regulatory activity and greater ASE, focusing on differential patterns of TF binding. Of the 421 sites, the majority (n= 239) had greater regulatory activity in the unmethylated condition compared to the methylated condition (as was observed in the overall dataset; binomial test, p=6.28×10^−3^). However, when examining MD genetic effects, we found that greater genetic effects on expression were equally likely in the methylated condition (n= 226) as the unmethylated condition (n=195; binomial test, p=0.144). The most common pattern suggested TF binding with limited regulatory activity in the methylated condition and allele specific binding in the unmethylated condition (29% of sites; Figure 4A). We also investigated whether the direction of ASE could vary between conditions (i.e., enhanced reference allele expression in one condition, enhanced alternate allele expression in the other). We found that 95 out of the 421 sites MD genetic effect sites not directly abolishing a CpG site have median reference allele expression greater than the DNA input in one condition, and less than the DNA input in the other condition (Figure 4B), suggesting mechanisms that reverse the direction of ASE depending on methylation status.

**Figure 4.**
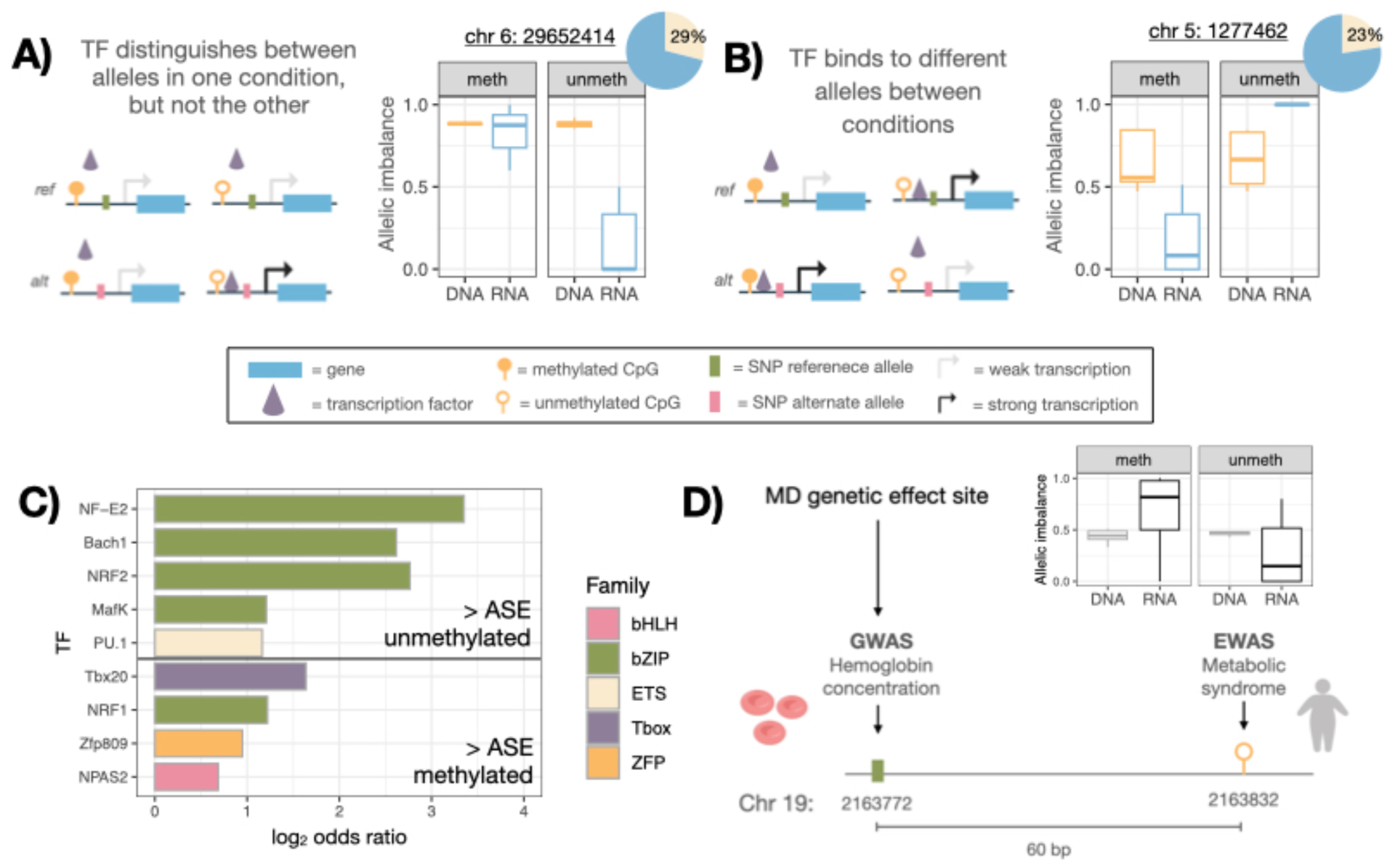
Insights into potential mechanisms involved in methylation-dependent genetic effects. A) One potential mechanism leading to MD genetic effects wherein TFs distinguish between alleles in one condition but not the other with an example site showing allelic imbalance (the ratio of reference allele to total counts) in the methylated versus unmethylated condition and a pie chart showing the percent of MD genetic effect sites following this pattern, B) another potential mechanism leading to MD genetic effects wherein TFs bind to different alleles in alternate conditions with an example site and a pie chart showing the percent of MD genetic effect sites following this pattern, C) TF motifs that are enriched within +/−200 bp of MD genetic sites that display increased ASE in the unmethylated condition (top) or methylated condition (bottom) colored by TF family, D) example of a MD genetic effect site that directly overlaps with a GWAS hit and is 60 bp away from an EWAS hit, which are associated with hemoglobin concentration and MetS, respectively. Created with Biorender.com

We next explored which TF binding motifs were enriched within +/− 200 bp of non-CpG altering MD genetic effect sites. Near sites that displayed greater ASE in the methylated condition, we found a significant enrichment for T-box binding motifs (10% FDR threshold) and a moderate enrichment binding motifs for TFs in the basic Leucine Zipper (bZIP), zinc-finger protein (ZFP), and basic Helix-Loop-Helix (bHLH) families (20% FDR threshold), relative to the background set of ASE sites that did not have MD effects. Conversely, near sites that displayed greater ASE in the unmethylated condition, we found a significant enrichment for bZIP and Erythroblast Transformation-Specific (ETS) binding motifs (10% FDR threshold; Figure 4C; Table S10). These two TF families have both been previously shown to preferentially bind to unmethylated sequences in mSTARR-seq (Lea et al. 2018) and other assays (Hernandez-Corchado and Najafabadi 2022). To assess in vivo TF binding, we used ChIP-Atlas, a database which integrates ChIP-seq peaks from thousands of experiments (Zou et al. 2024). Similar to our motif analysis, we compared either sites that displayed greater ASE in the methylated condition or sites that displayed greater ASE in the unmethylated condition to a set of matched background sites (see SI text for methods). We found 37 TFs that were more likely to be bound at variants associated with ASE in the methylated condition and 9 TFs that were more likely to be bound at variants associated with ASE in the unmethylated condition (Table S11). TFs associated with ASE in the methylated condition included 11 from the ZFP family, including CTCF, and 4 from the bZIP family, similar to our motif results.

Next, we overlapped MD genetic effect sites with both GWAS and EWAS datasets to better understand how both genetic and epigenetic variation may contribute to organism-level phenotype. If MD genetic effects occur at trait-associated sites identified in GWAS studies, DNAm might impact disease penetrance (i.e., the likelihood that a SNP contributes to a particular phenotype or disease). Likewise, MD genetic effect sites that overlap with trait-associated EWAS sites have the potential to increase our mechanistic understanding of how genetic and epigenetic factors contribute to phenotypes. We found that 98 MD genetic effect sites (17%) were located within 400 bp of a GWAS hit (75 of which directly overlapped with a GWAS hit), representing 151 different site-trait associations and 114 unique traits (Table S12). The most common traits associated with MD genetic effect sites included physical attributes like height and body mass, and other blood-related traits like white blood cell, red blood cell, and platelet count. When we asked whether MD genetic effect sites were associated with a greater number of GWAS hits or trait associations (as each GWAS hit can be associated with multiple traits) compared to random variants matched for gene density, LD, and MAF (see SI text for methods), we found that MD genetic effect sites were associated with a greater number of GWAS hits (permutation test: p=0.002; Figure S11), and a comparable number of trait associations (permutation test: p=0.3; Figure S11) compared to our matched control SNP sets. These results emphasize the utility of multiplexed mSTARR-seq in identifying variants that are important for organism-level phenotype.

Likewise, we found that 159 MD genetic effect sites (28%) were located within 400 bp of at least one EWAS probe, totaling 340 unique probes (EWAS utilizes probes to query specific CpG sites, as opposed to testing all CpG sites in the genome). Of these 340 probes, 134 were associated with one or more phenotypes (39.4%), resulting in 278 different site-trait associations (Table S13). Again, some traits were associated with multiple sites in our dataset, with the most common being aging and smoking, which were associated with 18 and 16 sites, respectively. To understand whether we observe more phenotype associations than would be expected by chance, we again utilized 1,000 matched control SNP datasets and tested whether MD genetic effect sites that were tested in the EWAS framework (i.e., were within 400 bp of an EWAS probe) were associated with a greater number of EWAS hits or trait associations. We found that MD genetic effect sites were not associated with more EWAS hits after controlling for the number of queried sites (permutation test: p=0.12; Figure S12), likely because EWAS testing does not query every CpG site in the genome, but instead focuses efforts in areas with likely functional importance (e.g., promoters, gene bodies, CpG islands). However, MD genetic effect sites were associated with a greater number of traits compared to the EWAS hits in our matched control SNP sets (permutation test: p= 0; Figure S12), suggesting that MD genetic effect sites contribute broadly to phenotypes, through pleiotropy or other mechanisms.

Finally, we find that 31 of our MD genetic effect sites are located within 400 bp of both a GWAS and an EWAS hit; these 31 sites are associated with 41 GWAS hits, 40 EWAS hits and 62 unique combinations of traits (Table S14). We found multiple GWAS trait-EWAS trait associations for related phenotypes; for example, a MD genetic effect site directly overlapped with a GWAS hit associated with hemoglobin concentration that is 60 bp away from an EWAS hit associated with metabolic syndrome (MetS; Figure 4D). Additionally, we found one example for which the GWAS and EWAS hit were associated with the same trait: we identified a MD genetic effect site that directly overlapped with a GWAS hit, and was 37 bp away from an EWAS hit, that were both associated with smoking.

### Multiplexing increases capacity to identify allele specific and methylation-dependent genetic effects

To gauge the potential benefits of multiplexing DNA from genetically diverse populations, we assessed both the number of SNPs located within regulatory regions and the number of MD genetic effect sites that could be identified in our dataset by subsampling different numbers of individuals included in the assay. We found that, on average, approximately 15x more unique SNPs can be analyzed when multiplexing 12 individuals as opposed to performing allele specific analyses on only a single individual (as has been done previously in STARR-seq experiments: Johnson et al. 2018). This is augmented especially by the inclusion of African individuals, with a nearly 20% increase in the number of SNPs found in regulatory regions when sampling individuals of African origin in comparison to European origin (Figure 5A). For example, when we limit the tested genomic regions to the 6,957 400 bp windows with significant regulatory function in either the methylated or unmethylated condition and subsample to sequences from a single European individual, we find that we can assay a mean of 65.4 (range: 19-123) variants. However, when we include sequences from all 12 of the European individuals in our assay, we find that we are able to assay 910 variants; these same numbers rise to 75.4 (range: 3-321) variants for a single African individual and 1104.9 (range: 1017-1205) for 12 multiplexed African individuals, respectively.

**Figure 5.**
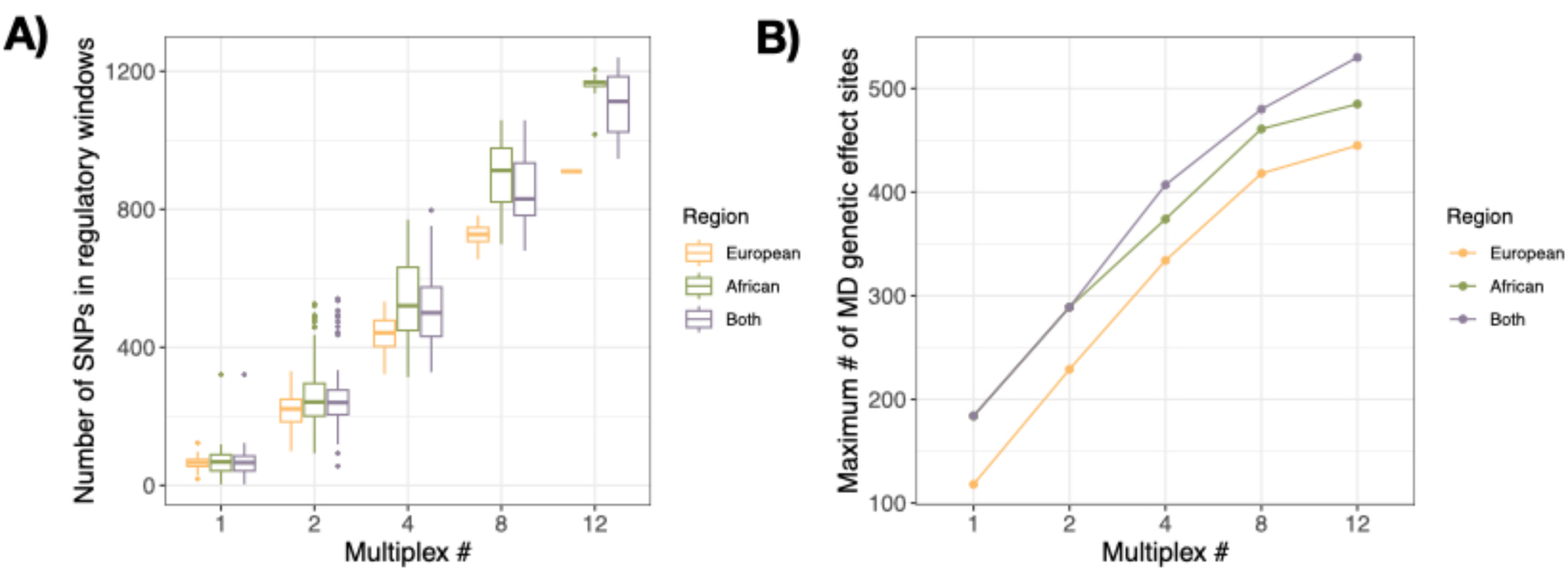
Multiplexing increases detection of methylation-dependent genetic effects. A) number of variant sites located in regulatory regions that could potentially be tested for ASE when subset for different multiplexing regimes and samples originating from different geographical regions, B) maximum number of MD genetic effect sites that would’ve been present in our assay when subset for different multiplexing regimes and geographical regions.

In addition to exploring allele specific effects on expression, we aimed to describe MD genetic effects, and thus we investigated how many of the 575 identified MD genetic effect sites would have been present and variable in our assay under different subsampling regimes. For example, sequences from a single European individual contained a maximum of 118 MD genetic effect sites and from a single African individual a maximum of 184 MD genetic effect sites. Notably, multiplexing 12 samples originating from both continents resulted in the greatest representation of the MD genetic effect sites identified in our experiment (530 sites; Figure 5B), arguing for inclusion of global genetic diversity whenever possible.

## DISCUSSION

Here, we present the first demonstration of multiplexed mSTARR-seq and its capacity to jointly assay the impact of methylation and genetic variation on regulatory element function. When identifying regulatory and MD regulatory regions, multiplexed mSTARR-seq performs similarly to previously, non-multiplexed iterations (Lea et al. 2018, Johnston et al. 2024), confirming that the method is repeatable and robust to the addition of genetic variation and multiplexing barcodes. New to this assay, however, we are able to assess how naturally occurring genetic variation sampled from 25 individuals from populations around the world, contributes to variation in regulatory activity and interacts with DNAm to have downstream impacts on gene expression. Through this, we identified several hundred instances where genotype modulated expression differently in methylated and unmethylated contexts.

eQTL studies have now identified variants associated with expression levels for nearly every gene in the genome (GTEx Consortium 2020), however, it is unclear how greatly these sites contribute to organism level phenotypes as many GWAS hits don’t colocalize with eQTL in any tissue (Umans et al. 2021). In a new study exploring the genomic contexts in which eQTL and GWAS hits are found, researchers observed that eQTL were more likely to be located within promoter regions close to TSSs, while phenotypically relevant GWAS hits were more likely to be depleted in promoter regions and located distally to TSSs (Mostafavi et al. 2023). Here, we find that in a causal, experimental assay, SNPs with allelic specific impacts on expression are found in similar genomic contexts to GWAS hits. They are located farther away from TSSs and are less likely to be located within promoter regions in comparison to SNPs with no impact on expression. Although GWAS studies are effective at linking genomic regions to traits, they often lack the ability to pinpoint causal variants among those in strong LD. While fine mapping can improve our ability to recognize causal variants among linked sites, MPRAs, including multiplexed mSTARR-seq, will be useful tools for experimentally nominating causal variants. For example, new efforts which utilize MPRAs to assay synthetic oligos (with a single variant per query fragment) have been effective in narrowing down 221,412 trait-associated fine mapped variants to 12,025 putatively causal variants that directly impact gene expression (Siraj et al. 2023). Notably, these assays (and similar previous work: Tewhey et al. 2016; Ulirsch et al. 2016) were conducted with all fragments in an entirely unmethylated state: our work suggests that an appreciable proportion of SNPs may have different effects on expression when nearby CpGs are methylated.

We explore several mechanisms that may generate MD genetic effects. First, many (26.8%) MD genetic effects involve a SNP that changes either the C or G base of a CpG site, thus directly inhibiting methylation for one genotype. This provides a straightforward mechanism by which methylation treatment can have no appreciable impact on gene expression for one genotype (because methylation cannot occur), while having a significant impact on expression for the other genotype. Second, TFs may exhibit genotype based binding preferences, and if the effects of methylation outweigh genotype preferences, these effects may only be observed in one condition. This pattern is evident in 29% of the MD genetic effect sites in our sample that had greater regulatory activity and a greater genetic effect in the unmethylated condition, suggesting that TF binding was inhibited in the methylated condition regardless of genotype. Likewise, 27.8% of MD genetic effects sites in our sample had greater regulatory activity in the unmethylated condition, but a greater genetic effect in the methylated condition, suggesting that TF affinity was so strong in the unmethylated condition that binding occurred regardless of genotype, a pattern that has also been observed previously (Kribelbauer et al. 2017). Third, TFs may preferentially bind to distinct sequences in methylated versus unmethylated contexts, a phenomenon that has been experimentally confirmed via electrophoretic mobility shift assays for 4 human TFs (Hu et al. 2013). In our assay, we found 95 sites in which allelic imbalance was in opposite directions in the methylated and unmethylated conditions, suggesting this process may extend to other TFs.

To understand which TFs may be particularly sensitive to variation in both sequence and methylation status, we investigated which TF binding motifs were proximal to MD genetic effect sites. Among sites with greater ASE in the unmethylated condition, we found three significantly overrepresented motifs: PU.1 from the ETS family and Bach-1 and NF-E2, both from the bZIP TF family. These TF families are both known to avoid binding to methylated sequences determined both from prior mSTARR-seq assays (Lea et al. 2018) and through joint accessibility-methylation-sequence modeling (Hernandez-Corchado and Najafabadi 2022). These results were corroborated by ChIP-seq data, wherein we find NF-E2, as well as 2 other TFs within the bZIP family (FOSL1 and ATF2), are more likely to be bound at sites exhibiting ASE in the unmethylated condition compared to matched background sites. Among sites with greater ASE in the methylated condition, we found a significant enrichment for Tbx20 binding motifs. Tbx20 is a TF known to play a pivotal role in heart development and is a part of the Box family proteins, which have been shown to preferentially bind to methylated sequences in other studies (Hu et al. 2013). Just above our significance threshold, both NRF1 and Zfp809 (FDR adjusted p-values = 0.18) were also enriched near sites with greater ASE in methylated condition. NRF1 is part of the bZIP TF family, which has been demonstrated via SELEX binding assays to be particularly sensitive to the position of CpG methylation in the flanking regions of binding motifs (Kribelbauer et al. 2017). Zfp809 is a KRAB ZFP, a family of TFs containing a canonical methyl-binding domain (Hashimoto et al. 2015). The ability of ZFPs to engage in allele specific binding in methylated contexts is further bolstered by ChIP-seq data, wherein 30% of the TFs found to be enriched in ASE in the methylated condition were members of the ZFP family. Prior MPRA studies have also found an enrichment for these same TF families near to areas of the genome containing multiple variants in strong LD associated with eQTL and GWAS hits (Abell et al. 2022).

MD genetic effect sites are enriched for GWAS hits and EWAS trait-associations, suggesting that these sites are particularly relevant for downstream phenotypes. Further, there are several examples within our dataset suggesting that the GWAS and EWAS hits associated with MD genetic effect sites may be associated with the same underlying condition. For example, we found a MD genetic effect site directly overlapping with a GWAS hit for hemoglobin concentration (chr19: 2163772) and in close proximity to an EWAS site associated with metabolic syndrome (MetS; chr19: 2163832). Multiple publications have documented elevated hemoglobin within patients with MetS, suggesting that these may be part of the same etiology (Wang et al. 2004; Lohsoonthorn et al. 2007; Nebeck et al. 2012). For this site, we observe regulatory activity in both conditions but with different alleles preferentially expressed in the methylated versus unmethylated condition (Figure 4E). Furthermore, our estimation of the proportion of GWAS sites that co-occur with an EWAS site and exhibit regulatory activity in our assay is likely underestimated, as variant sites may be sensitive to methylation only in certain tissues or cellular context (Umans et al. 2021).

By developing a multiplexed design, we are able to assay genetic variation from 25 individuals for regulatory and MD regulatory effects in a single experiment, allowing us to test a much greater number of variants for both ASE and MD genetic effects than previous work (Lea et al. 2018, Johnston et al. 2024). An over-representation of European ancestry individuals is a current reality of most ‘omics’ datasets, with >80% of GWAS and EWAS participants reporting European ancestry (Mills and Rahal 2020; Breeze et al. 2022). However, the inclusion of non-European individuals in association studies improves not only our ability to identify disease-related variants specific to marginalized populations, but also improves statistical power to identify disease-relevant variants across *all* populations (Pulit et al. 2010). Thus, the development of new technologies which support the inclusion of diverse genetic material into experimental assays will be necessary to develop a more comprehensive understanding of the complex interactions between genetic variants, the epigenome, and gene expression. Moving forward, the incorporation of additional aspects of variation into MPRAs, including new cell types and cellular environments, as well as the expansion to non-human species (see Hansen et al. 2024), will provide important new biomedical and evolutionary insight into the role of gene regulation in the generation of complex traits.

## METHODS

### Input library generation

We generated input fragments by extracting genomic DNA from lymphoblastoid cell lines from 25 individuals included in the 1000 Genomes Project. The cell lines were obtained from Coriell Institute for Medical Research. We included 12 individuals of European origin and 13 individuals of African origin, sampled from 5 different locations each (Table S1). For each sample, we performed an *Msp1* digest using 3ug of genomic DNA, in a 25ul volume incubated overnight at 37°C. *Msp1* has a CCGG recognition motif and thus enriches for ∼5% of the genome containing a high density of CpG sites (Gu et al. 2011). We size selected the resultant Msp1 digested fragments by gel electrophoresis and the Zymoclean Gel DNA Recovery kit, focusing on the 300-700 bp range. We ligated an adapter to each size-selected DNA fragment pool using the NEB Ultra II DNA Library Prep Kit following the manufacturer’s instructions for end prep, adaptor ligation, and size selection for a 400-500 bp insert size distribution in half volume reactions. We performed PCR enrichment of adaptor ligated DNA using 25ul of NEB Q5 hotstart reaction mix, 15ul of size selected adaptor ligated DNA fragments, 2.5ul of 10 umol mSTARR F2 primer, 2.5 ul of mSTARR R2 primer (with the new addition of an individual-specific, CpG free barcode) and 5ul of water with the following cycling conditions: initial denaturation at 98°C for 30 seconds, 12 cycles of denaturation at 98°C for 10 seconds, annealing at 62°C for 30 seconds, extension at 72°C for 60 seconds, and a final extension at 72°C for 5 minutes. All relevant mSTARR primer sequences can be found in Table S2.

We created two pools of barcoded libraries that were used as input for our experiments and derived from independent library preparations. Pool 1 contained 20 individuals and pool 2 contained 21 individuals (Table S3). 16 individuals are represented in both pools and 9 individuals are included in only one of the two pools. When a sample was included in both pools, we used different multiplexing indices in pool 1 versus pool 2 to ensure that results were reproducible across independent library preparations. We also included a subset of samples twice within the same pool with two different indices (n=2 in each pool). By sequencing most individuals twice with different indices, our goal was to limit any potential impacts that the barcode sequence we included to differentiate individual libraries may have on our results.

### mSTARR-seq

The resulting pools of barcoded libraries were used as input into the mSTARR-seq assay, closely following the protocol published in Lea et al. 2018. Briefly, we linearized the mSTARR plasmid and inserted the multiplexed *MspI*-digested libraries using Gibson assembly. To replicate the assembled plasmid libraries, we transformed each pool into electrocompetent GT115 cells, incubated overnight, and extracted the replicated plasmid libraries. We then exposed half of each plasmid pool to water (mock/sham control) and the other half to a CpG methyltransferase enzyme (*M.SssI*) to experimentally induce methylation. We transfected the treated plasmids into K562 cells, performing 3 replicates of each condition (3 methylated and 3 unmethylated replicates) for each of the 2 pools, creating 12 replicates total with 15 million cells each. We incubated the transfected cells for 48 hours before performing DNA/RNA extraction. We extracted DNA from approximately 3 million cells and extracted RNA from the remaining approximately 12 million cells from each of the 12 replicates. We created mRNA sequencing libraries by first isolating mRNA and then performing a targeted cDNA synthesis using a custom primer specific to the mSTARR plasmid (RT_polyA; Table S2). We then amplified the resultant cDNA using a nested, two-step PCR which first amplifies each pool using a targeted primer that spans the splice junction of the synthetic intron contained in the mSTARR plasmid (PCR1_F_splice; Table S2), followed by a second PCR to add NEBNext Multiplex Oligos for Illumina. We quantified our final DNA and mRNA libraries, pooled, and sequenced on the NovaSeq6000 platform using a single S4 flowcell and 100 bp PE reads. A more detailed description of how we performed the mSTARR-seq protocol can be found in the SI text, and read counts and summary stats per sample are available in Table S3.

### Data processing

Following sequencing, we trimmed adapter sequences and low quality reads using cutadapt v 1.16 (Martin 2011). We used BWA v 0.7.17 to map the trimmed reads to the chromosomal assemblies of the *hg38* human reference genome and kept only uniquely mapped reads (Li and Durbin 2009). A small percentage of the total reads generated for each sample mapped to the Eppstein-Barr virus genome, the method used to immortalize B lymphocytes into LCL cell lines (mean: 4.5%, SD: 3.5%). We determined the number of unique input DNA fragments per replicate based on the number of reads with unique start and end mapping locations. To determine the precision of our Msp1 digested input library, we compared the fragments in our input library to those that would be expected by performing in silico digest of the human genome (*hg38*), cutting (bioinformatically) at all instances of CCGG, and retaining fragments between 100-700 bp. We then mapped the in silico generated fragments, converted the mapped reads into a bed file with genomic locations, and used the bedtools ‘intersect’ function (Quinlan and Hall 2010) to determine the proportion of in silico digest fragments that were covered in our input library with a minimum overlap of 75% of the fragment length.

To determine regulatory and methylation-dependent regulatory activity, averaged across genotypes, we combined uniquely mapped reads from all individuals within a replicate (n=11 replicates) and overlapped these with 400 bp non-overlapping genomic windows, as this was the approximate size of our inserts. We calculated coverage of each window using the bedtools ‘coverage’ function and filtered for windows based on coverage of both DNA and RNA. We retained windows that possessed at least one DNA read in at least half of the replicates in both conditions (i.e., ≥ 1 read in 3 or more of the methylated replicates *and* ≥ 1 read in 3 or more of the unmethylated replicates) and at least one RNA read in at least half of the replicates in either condition (i.e., ≥ 1 read in 3 or more of the methylated replicates *or* ≥ 1 read in 3 or more of the unmethylated replicates). This filtering left us with 525,074 400 bp windows. We determined the number of CpG sites per window using the bedtools ‘intersect’ function and a bed file of all CpG sites in the human genome (*hg38*). We also explored the location of these windows by overlapping them with annotations for gene bodies, promoters and CpG islands. We gathered the locations of CpG islands from the UCSC Table Browser CpG islands track and gene body locations from Ensembl 108 with promoters defined as the 10kb region upstream of each gene.

### Identification of regulatory and methylation-dependent regulatory elements

We used linear mixed effects modeling to test for regulatory activity at each 400bp window in methylated and unmethylated conditions separately. We normalized total read counts for the 11 DNA and 11 RNA replicates using ‘voomWithQualityWeights’, fit a linear model for each window within each condition using ‘lmFit’, and calculated test statistics using ‘eBayes”, all of which are functions available in the R package ‘limma’ (Ritchie et al. 2015). We included sample type (DNA vs RNA) as our predictor variable, pool (1 vs 2) as a covariate, and normalized counts as the response variable in each linear model. Using the p-value output for each condition, we extracted p-values for the sample type term and performed a FDR correction using the R function ‘qvalue’ (Storey et al. 2024). We considered a region to have regulatory activity in a given condition if 1) sample type (DNA vs RNA) was a significant predictor of read count at an FDR corrected p-value < 0.01, and 2) the effect size for sample type was > 0, meaning that the normalized counts of mRNA were greater than the normalized counts of DNA for that window. This left us with 6,221 and 2,513 regulatory regions in the unmethylated and methylated conditions, respectively.

To both validate our results and generate a better understanding of the genetic contexts in which mSTARR-seq identifies regulatory activity, we ascertained which chromatin state regulatory windows are located in by overlapping significant windows with chromatin state annotations generated for K562 cells, lifted over from the hg19 to hg38 genome build (ENCODE Project Consortium 2012; Hoffman et al. 2013). We ran a Fisher’s exact test to quantify enrichment of different chromatin states within our regulatory windows (Table S6). We also confirmed our results by reprocessing two previous mSTARR-seq datasets using the same data analysis pipeline outlined above, and tested for a correlation in effect sizes and an enrichment in shared regulatory activity (see SI text for more detailed description).

We estimated methylation-dependent regulatory activity within our dataset using the ‘mashr’ package in R (Urbut et al. 2019). This package provides tools to generate more accurate effect size estimates across multi-condition datasets by exploiting correlations between conditions; it also provides a framework for testing for effect size sharing versus heterogeneity across conditions. Using the results from our linear modeling, we followed the authors’ recommendations and input the 6,957 regions that met our regulatory activity criteria in at least one condition as the set of ‘strong sites’ which mashR uses to learn the correlation structure between non-null effects in different conditions; we also included a random set of windows (n=20,000) to estimate signals in the data associated with null effects, and distinguish these from non-null effects. After running mashR with this pipeline, we generated a posterior mean and LFSR for each of our set of 6,957 tested regions. We subtracted the posterior mean of the sample type effect estimated in the methylated condition from the unmethylated condition to generate a logFC estimate, which represents the difference in regulatory activity between conditions. We considered a window to have MD activity if the LFSR was < 0.1 and the logFC between conditions was > 1.5. To determine if CpG density was associated with methylation-dependence, we overlapped MD windows with all CpG sites in the human genome and ran a T-test comparing the number of CpG sites found in MD versus non-MD windows.

### Data processing for allele specific expression

To interrogate the effect of genotype on regulatory activity, we estimated allele specific RNA output versus DNA input in the methylated and unmethylated conditions separately. We merged mapped DNA bam files for each individual within each pool and performed SNP calling for each individual (n=25) using GATK v 4.1.4.0 and following GATK’s best practices guidelines (Van der Auwera et al. 2013). From these SNP calls, we performed joint genotyping using the GenotypeGVFs function and generated allele specific read counts using the ‘ASEReadCounter’ function, both from the GATK package. We performed numerous filtering steps (detailed in the SI text) which limited our final SNP set to: 1) high confidence biallelic SNPs with a MAF within the 1000 Genomes dataset of greater than 0.05, 2) SNPs with regulatory activity in our window-based analysis that were not closely linked (within 400 bp) with another SNP, and 3) SNPs with adequate coverage and allelic variation required for modeling. These steps left us with a final number of 3,037 variants in the methylated condition and 4,931 variants in the unmethylated condition. The number of retained SNPs following each filtering criteria can be found in Table S5.

### Analysis of allele specific expression data

For the 3,037 sites in the methylated condition and 4,931 sites in the unmethylated condition that passed our filtering criteria, we ran beta binomial models using the function ‘betabin’ in the ‘aod’ package in R to test whether the ratio of reference to alternate allele counts significantly differed between the DNA input and the mRNA output at each site (Lesnoff and Lancelot 2012). We included sample type (DNA vs RNA) as the predictor variable, pool as a covariate, and the number of reference and alternate allele counts as the response. Similar to our initial analyses of regulatory activity pooled across genotypes, we ran these analyses within each condition separately. We explored the genomic context of genetic effect sites utilizing a dataset containing both sites with ASE and mono/ nearly monoallelic sites (although results are robust to the inclusion of only ASE sites: see SI text). We ascertained chromatin states for these sites by overlapping with the same chromatin state annotations for K562 cells as mentioned above and performed a Fisher’s exact test to assess the enrichment of enhancer states.

There are distinct differences in where GWAS versus eQTL hits are found, which may be related to the phenotypic importance of these sites. GWAS hits are linked to organismal level traits, and are located farther from TSSs and are depleted in promoter regions in comparison to a background set of variable genomic sites (Mostafavi et al. 2023). Conversely, eQTL sites are more commonly located within promoter regions closer to TSSs, however their phenotypic relevance is oftentimes unclear (Umans et al. 2021; Connally et al. 2022). To explore whether genetic effect sites identified using mSTARR-seq are more comparable to GWAS or eQTL hits based on their genomic location, we utilized the dataset containing both ASE and mono/ nealy monoallelic sites and summarized the distance to the nearest transcription start site (TSS) and promoter annotations for each site using the ‘annotatePeaks’ function in HOMER v 4.11.1 (Heinz et al. 2010). We performed a T-test to compare the distance to the nearest TSS for ASE sites versus the matched background set of variable sites and a Fisher’s exact test to examine under-enrichment of promoter annotations in ASE sites compared to the matched background.

### Analysis of methylation-dependent genetic effects

To understand how methylation impacts allele specific expression, we again used mashr to compare effect size estimates between conditions. In this case, we are asking whether the genetic effect on regulatory activity is modified by experimental methylation. We tested the 1,359 sites that had adequate coverage/ variability between replicates to be modeled in both conditions (i.e., were not removed due to mono/ nearly mono-allelic expression). We input 783 sites that had strong ASE (at a 1% FDR) in at least one condition as the set of ‘strong sites’, and included a random set of 783 sites as those with null effects to generate a posterior mean and LFSR for each ‘strong site’. Beta binomial modeling uses raw reference and alternate allele counts that are not normalized across libraries, so we log2 transformed the posterior mean of the unmethylated and methylated conditions before calculating the logFC between conditions (unmethylated condition divided by the methylated condition). We considered a SNP to have methylation-dependent genetic effects if the logFC between conditions was > 1.5.

One potential mechanism by which MD genetic effects might arise is due to variable binding in TFs. To better understand how methylation may impact allele specific TF binding, we used the ‘findMotifsGenome’ function in the program HOMER to identify transcription factor binding motifs enriched within a +/− 200 bp range centered around the variant site of interest (Heinz et al. 2010). We split our MD genetic effect sites into two sets: those with a more pronounced genetic effect in the methylated condition (n= 226) and those with a more pronounced genetic effect in the unmethylated condition (n= 195). We used sites with ASE whose effects were not modified by methylation status as the background comparison set (methylated condition: n= 776, unmethylated condition: n= 780). P-values for enrichment of TF binding motifs were adjusted for multiple hypothesis testing using the Benjamini-Hochberg correction (Benjamini and Hochberg 1995). Additionally, we assessed whether the abundance of TF binding motifs translated to differences in TF binding by incorporating ChIP-seq peaks curated by ChIP-Atlas 3.0 (Zou et al. 2024). Using this program, we again asked whether there was enrichment for TF binding among sites with a more pronounced genetic effect in the methylated condition or unmethylated condition compared to a background set of SNPs that were matched for MAF, LD, and gene density. Further information on how we performed SNP matching to create the background set can be found in the SI text. We also required that SNPs included in our background were ones that could have potentially been included in our assay by filtering for SNPs within the 400 bp windows that possessed at least one DNA read in at least half of the replicates in both conditions of our assay, so as not to bias our results towards TFs which bind to GC-rich regions. We set our threshold of significance to include peaks with a q-value of less than 1×10^−50^ (generated using MACS2 peak-caller), filtered results for K-562 cells, and report enriched TFs at a 1% FDR.

We further explored the phenotypic relevance of MD genetic effect sites by intersecting these sites with known GWAS and EWAS hits, using the GWAS catalog v1.0.2 associations (Sollis et al. 2023) and EWAS atlas associations accessed through the EWAS open platform data hub (Li et al. 2019), and downloaded in March 2024. We identified the number of MD genetic effect sites located near trait-associated sites by searching within ± 400 bp of each GWAS hit and EWAS probe. To understand how the number of GWAS and EWAS associations found in our dataset compares to what would be expected by chance, we compared our results to the number of associations that would be found using 1000 datasets of SNPs that have been matched for MAF, LD, and gene density. Further information on how these matched datasets were created can be found in the SI text. Similarly, we identified the number of sites within each of the 1000 matched datasets that were located within ± 400 bp of a GWAS hit or EWAS probe. To determine whether MD genetic effect sites are enriched in GWAS or EWAS hits, or are associated with a greater number of traits (as each GWAS and EWAS hits can be associated with multiple traits), we performed a permutation test and report the p-value as the proportion of matched sets having a greater number of associations compared to our test set (MD genetic effect sites).

### Assessing the benefits of multiplexing

To determine how the multiplexing of genetically diverse input fragments impacts our capacity to identify MD genetic effects, we performed two analyses. First, we determined the number of SNPs that would be identified in regulatory fragments when down sampling our dataset to combinations of 1, 2, 4, 8, or 12 individuals. We intersected the joint genotyping vcf containing the genotype found for each library at each variable site with the locations of our 6,957 regulatory windows identified in either condition, then downsampled the number of libraries included in our calculations to mimic different potential multiplexing regimes. Since some individuals had libraries in both pools or two libraries in the same pool, we limited our analyses to include only one library per individual, keeping each individual’s library with the highest mean DNA read count across replicates. When downsampling to a single individual, we extracted the number of heterozygous sites located within a regulatory window for each of the 25 individuals. When downsampling to 2, 4, 8, or 12 individuals, we created 100 unique combinations of individuals (or the maximum unique combinations possible) for each multiplexing regime and counted the number of variable sites that would have had both the reference and alternate allele present in the assay. To better understand how geographic origin may impact the amount of genetic diversity assayed, we ran this analysis 3 times, using genotype information from all European sampled individuals, all African sampled individuals, or both.

Second, we compared the maximum number of MD genetic effect sites identified in the current study that would have been included in the assay within different multiplexing regimes. To do so, we intersected the joint genotyping vcf with the MD genetic effect sites, and summed the number of unique MD genetic effect sites that would have had both alleles represented in the assay when downsampling to 1, 2, 4, 8, or 12 individuals. Again, we limited this analysis to include one library from each individual and created a distribution of the number of MD genetic effect sites present by sampling 100 unique combinations of individuals to include in each multiplex regime. We performed this analysis 3 times, subsetting to individuals of European origin, African origin, or all individuals.

## DATA ACCESS

The DNA and RNA sequencing reads generated in this study have been submitted to the National Center Biotechnology Information Short Read Archive (https://www.ncbi.nlm.nih.gov/sra) under accession number PRJNA1137064. Counts matrices, modeling results, and other datasets used in the text are available on Zenodo, DOI: 10.5281/zenodo.12763233. Code is available at https://github.com/rachpetersen/muliplexed_mSTARRseq.

## ACKNOWLEDGMENTS

We would like to thank members of the Lea lab, Hodges lab, and the Evolutionary Studies Initiative at Vanderbilt University for their support in completing this work. We specifically thank Emily Hodges, Tyler Hansen, Sai Han Presley, Nicholas Ryan, Rachel Johnston, and Jenny Tung for their advice and input. This work was supported by the Searle Scholars Program and NIH/NIGMS (1R35GM147267).

## REFERENCES

Abell NS, DeGorter MK, Gloudemans MJ, Greenwald E, Smith KS, He Z, Montgomery SB. 2022. Multiple causal variants underlie genetic associations in humans. Science 375: 1247–1254.

Anderson JA, Lin D, Lea AJ, Johnston RA, Voyles T, Akinyi MY, Archie EA, Alberts SC, Tung J. 2024. DNA methylation signatures of early-life adversity are exposure-dependent in wild baboons. Proceedings of the National Academy of Sciences 121: e2309469121.

Arnold CD, Gerlach D, Stelzer C, Boryń ŁM, Rath M, Stark A. 2013. Genome-wide quantitative enhancer activity maps identified by STARR-seq. Science 339: 1074–1077.

Banovich NE, Lan X, McVicker G, Van de Geijn B, Degner JF, Blischak JD, Roux J, Pritchard JK, Gilad Y. 2014. Methylation QTLs are associated with coordinated changes in transcription factor binding, histone modifications, and gene expression levels. PLoS genetics 10: e1004663.

Benjamini Y, Hochberg Y. 1995. Controlling the false discovery rate: a practical and powerful approach to multiple testing. Journal of the royal statistical society Series B (Methodological): 289–300.

Breeze CE, Beck S, Berndt SI, Franceschini N. 2022. The missing diversity in human epigenomic studies. Nature genetics 54: 737–739.

Chun S, Casparino A, Patsopoulos NA, Croteau-Chonka DC, Raby BA, De Jager PL, Sunyaev SR, Cotsapas C. 2017. Limited statistical evidence for shared genetic effects of eQTLs and autoimmune-disease-associated loci in three major immune-cell types. Nature genetics 49: 600–605.

Connally NJ, Nazeen S, Lee D, Shi H, Stamatoyannopoulos J, Chun S, Cotsapas C, Cassa CA, Sunyaev SR. 2022. The missing link between genetic association and regulatory function. Elife 11: e74970.

Deplancke B, Alpern D, Gardeux V. 2016. The genetics of transcription factor DNA binding variation. Cell 166: 538–554.

ENCODE Project Consortium. 2012. An integrated encyclopedia of DNA elements in the human genome. Nature 489: 57.

Garg P, Joshi RS, Watson C, Sharp AJ. 2018. A survey of inter-individual variation in DNA methylation identifies environmentally responsive co-regulated networks of epigenetic variation in the human genome. PLoS genetics 14: e1007707.

Gong J, Mei S, Liu C, Xiang Y, Ye Y, Zhang Z, Feng J, Liu R, Diao L, Guo A-Y. 2018. PancanQTL: systematic identification of cis-eQTLs and trans-eQTLs in 33 cancer types. Nucleic acids research 46: D971–D976.

Greenberg MV, Bourc’his D. 2019. The diverse roles of DNA methylation in mammalian development and disease. Nature reviews Molecular cell biology 20: 590–607.

Griesemer D, Xue JR, Reilly SK, Ulirsch JC, Kukreja K, Davis JR, Kanai M, Yang DK, Butts JC, Guney MH. 2021. Genome-wide functional screen of 3′ UTR variants uncovers causal variants for human disease and evolution. Cell 184: 5247–5260. e5219.

GTEx Consortium. 2020. The GTEx Consortium atlas of genetic regulatory effects across human tissues. Science 369: 1318–1330.

Gu H, Smith ZD, Bock C, Boyle P, Gnirke A, Meissner A. 2011. Preparation of reduced representation bisulfite sequencing libraries for genome-scale DNA methylation profiling. Nature protocols 6: 468–481.

Gurdasani D, Barroso I, Zeggini E, Sandhu MS. 2019. Genomics of disease risk in globally diverse populations. Nature Reviews Genetics 20: 520–535.

Han H, Cortez CC, Yang X, Nichols PW, Jones PA, Liang G. 2011. DNA methylation directly silences genes with non-CpG island promoters and establishes a nucleosome occupied promoter. Human Molecular Genetics 20: 4299–4310.

Hansen TJ, Fong SL, Day JK, Capra JA, Hodges E. 2024. Human gene regulatory evolution is driven by the divergence of regulatory element function in both cis and trans. Cell Genomics 4.

Hashimoto H, Zhang X, Vertino PM, Cheng X. 2015. The mechanisms of generation, recognition, and erasure of DNA 5-methylcytosine and thymine oxidations. Journal of biological chemistry 290: 20723–20733.

Heinz S, Benner C, Spann N, Bertolino E, Lin YC, Laslo P, Cheng JX, Murre C, Singh H, Glass CK. 2010. Simple combinations of lineage-determining transcription factors prime cis-regulatory elements required for macrophage and B cell identities. Molecular cell 38: 576–589.

Hernandez-Corchado A, Najafabadi HS. 2022. Toward a base-resolution panorama of the in vivo impact of cytosine methylation on transcription factor binding. Genome Biology 23: 151.

Hoffman MM, Ernst J, Wilder SP, Kundaje A, Harris RS, Libbrecht M, Giardine B, Ellenbogen PM, Bilmes JA, Birney E. 2013. Integrative annotation of chromatin elements from ENCODE data. Nucleic acids research 41: 827–841.

Holliday R, Pugh JE. 1975. DNA Modification Mechanisms and Gene Activity During Development: Developmental clocks may depend on the enzymic modification of specific bases in repeated DNA sequences. Science 187: 226–232.

Hu S, Wan J, Su Y, Song Q, Zeng Y, Nguyen HN, Shin J, Cox E, Rho HS, Woodard C. 2013. DNA methylation presents distinct binding sites for human transcription factors. elife 2: e00726.

Inoue F, Ahituv N. 2015. Decoding enhancers using massively parallel reporter assays. Genomics 106: 159–164.

Jagoda E, Xue JR, Reilly SK, Dannemann M, Racimo F, Huerta-Sanchez E, Sankararaman S, Kelso J, Pagani L, Sabeti PC. 2022. Detection of Neanderthal adaptively introgressed genetic variants that modulate reporter gene expression in human immune cells. Molecular Biology and Evolution 39: msab304.

Jin Z, Liu Y. 2018. DNA methylation in human diseases. Genes & diseases 5: 1–8.

Johnson GD, Barrera A, McDowell IC, D’Ippolito AM, Majoros WH, Vockley CM, Wang X, Allen AS, Reddy TE. 2018. Human genome-wide measurement of drug-responsive regulatory activity. Nature communications 9: 5317.

Johnston RA, Aracena KA, Barreiro LB, Lea AJ, Tung J. 2024. DNA methylation-environment interactions in the human genome. Elife 12: RP89371.

Kalita CA, Moyerbrailean GA, Brown C, Wen X, Luca F, Pique-Regi R. 2018. QuASAR-MPRA: accurate allele-specific analysis for massively parallel reporter assays. Bioinformatics 34: 787–794.

Kaluscha S, Domcke S, Wirbelauer C, Stadler MB, Durdu S, Burger L, Schübeler D. 2022. Evidence that direct inhibition of transcription factor binding is the prevailing mode of gene and repeat repression by DNA methylation. Nature genetics 54: 1895–1906.

Kasowski M, Grubert F, Heffelfinger C, Hariharan M, Asabere A, Waszak SM, Habegger L, Rozowsky J, Shi M, Urban AE. 2010. Variation in transcription factor binding among humans. science 328: 232–235.

Kerimov N, Hayhurst JD, Peikova K, Manning JR, Walter P, Kolberg L, Samoviča M, Sakthivel MP, Kuzmin I, Trevanion SJ. 2021. A compendium of uniformly processed human gene expression and splicing quantitative trait loci. Nature genetics 53: 1290–1299.

Kim-Hellmuth S, Aguet F, Oliva M, Muñoz-Aguirre M, Kasela S, Wucher V, Castel SE, Hamel AR, Viñuela A, Roberts AL. 2020. Cell type–specific genetic regulation of gene expression across human tissues. Science 369: eaaz8528.

Kreibich E, Kleinendorst R, Barzaghi G, Kaspar S, Krebs AR. 2023. Single-molecule footprinting identifies context-dependent regulation of enhancers by DNA methylation. Molecular Cell 83: 787–802. e789.

Kribelbauer JF, Laptenko O, Chen S, Martini GD, Freed-Pastor WA, Prives C, Mann RS, Bussemaker HJ. 2017. Quantitative analysis of the DNA methylation sensitivity of transcription factor complexes. Cell reports 19: 2383–2395.

Kribelbauer JF, Lu X-J, Rohs R, Mann RS, Bussemaker HJ. 2020. Toward a mechanistic understanding of DNA methylation readout by transcription factors. Journal of molecular biology 432: 1801–1815.

Kwasnieski JC, Mogno I, Myers CA, Corbo JC, Cohen BA. 2012. Complex effects of nucleotide variants in a mammalian cis-regulatory element. Proceedings of the National Academy of Sciences 109: 19498–19503.

Lea AJ, Vockley CM, Johnston RA, Del Carpio CA, Barreiro LB, Reddy TE, Tung J. 2018. Genome-wide quantification of the effects of DNA methylation on human gene regulation. Elife 7: e37513.

Lesnoff M, Lancelot R. 2012. aod: Analysis of Overdispersed Data. R package version 1.3, URL.

Li H, Durbin R. 2009. Fast and accurate short read alignment with Burrows–Wheeler transform. bioinformatics 25: 1754–1760.

Li M, Zou D, Li Z, Gao R, Sang J, Zhang Y, Li R, Xia L, Zhang T, Niu G. 2019. EWAS Atlas: a curated knowledgebase of epigenome-wide association studies. Nucleic acids research 47: D983–D988.

Lohsoonthorn V, Jiamjarasrungsi W, Williams MA. 2007. Association of hematological parameters with clustered components of metabolic syndrome among professional and office workers in Bangkok, Thailand. Diabetes & Metabolic Syndrome: Clinical Research & Reviews 1: 143–149.

Loyfer N, Magenheim J, Peretz A, Cann G, Bredno J, Klochendler A, Fox-Fisher I, Shabi-Porat S, Hecht M, Pelet T. 2023. A DNA methylation atlas of normal human cell types. Nature 613: 355–364.

Luo C, Hajkova P, Ecker JR. 2018. Dynamic DNA methylation: In the right place at the right time. Science 361: 1336–1340.

Luo X, Zhang T, Zhai Y, Wang F, Zhang S, Wang G. 2021. Effects of DNA methylation on TFs in human embryonic stem cells. Frontiers in genetics 12: 639461.

Martin M. 2011. Cutadapt removes adapter sequences from high-throughput sequencing reads. EMBnet journal 17: 10–12.

Maurano MT, Humbert R, Rynes E, Thurman RE, Haugen E, Wang H, Reynolds AP, Sandstrom R, Qu H, Brody J. 2012. Systematic localization of common disease-associated variation in regulatory DNA. Science 337: 1190–1195.

Melnikov A, Murugan A, Zhang X, Tesileanu T, Wang L, Rogov P, Feizi S, Gnirke A, Callan Jr CG, Kinney JB. 2012. Systematic dissection and optimization of inducible enhancers in human cells using a massively parallel reporter assay. Nature biotechnology 30: 271–277.

Mills MC, Rahal C. 2020. The GWAS Diversity Monitor tracks diversity by disease in real time. Nature genetics 52: 242–243.

Monteagudo-Sánchez A, Noordermeer D, Greenberg MV. 2024. The impact of DNA methylation on CTCF-mediated 3D genome organization. Nature Structural & Molecular Biology 31: 404–412.

Mostafavi H, Spence JP, Naqvi S, Pritchard JK. 2023. Systematic differences in discovery of genetic effects on gene expression and complex traits. Nature Genetics 55: 1866–1875.

Nebeck K, Gelaye B, Lemma S, Berhane Y, Bekele T, Khali A, Haddis Y, Williams MA. 2012. Hematological parameters and metabolic syndrome: findings from an occupational cohort in Ethiopia. Diabetes & Metabolic Syndrome: Clinical Research & Reviews 6: 22–27.

Pacis A, Tailleux L, Morin AM, Lambourne J, MacIsaac JL, Yotova V, Dumaine A, Danckaert A, Luca F, Grenier J-C. 2015. Bacterial infection remodels the DNA methylation landscape of human dendritic cells. Genome research 25: 1801–1811.

Patwardhan RP, Hiatt JB, Witten DM, Kim MJ, Smith RP, May D, Lee C, Andrie JM, Lee S-I, Cooper GM. 2012. Massively parallel functional dissection of mammalian enhancers in vivo. Nature biotechnology 30: 265–270.

Patwardhan RP, Lee C, Litvin O, Young DL, Pe’er D, Shendure J. 2009. High-resolution analysis of DNA regulatory elements by synthetic saturation mutagenesis. Nature biotechnology 27: 1173–1175.

Perera BP, Faulk C, Svoboda LK, Goodrich JM, Dolinoy DC. 2020. The role of environmental exposures and the epigenome in health and disease. Environmental and Molecular Mutagenesis 61: 176–192.

Pulit SL, Voight BF, de Bakker PI. 2010. Multiethnic genetic association studies improve power for locus discovery. PloS one 5: e12600.

Quinlan AR, Hall IM. 2010. BEDTools: a flexible suite of utilities for comparing genomic features. Bioinformatics 26: 841–842.

Riggs AD. 1975. X inactivation, differentiation, and DNA methylation. Cytogenetic and Genome Research 14: 9–25.

Ritchie ME, Phipson B, Wu D, Hu Y, Law CW, Shi W, Smyth GK. 2015. limma powers differential expression analyses for RNA-sequencing and microarray studies. Nucleic acids research 43: e47–e47.

Salameh Y, Bejaoui Y, El Hajj N. 2020. DNA methylation biomarkers in aging and age-related diseases. Frontiers in Genetics 11: 480672.

Sharon E, Kalma Y, Sharp A, Raveh-Sadka T, Levo M, Zeevi D, Keren L, Yakhini Z, Weinberger A, Segal E. 2012. Inferring gene regulatory logic from high-throughput measurements of thousands of systematically designed promoters. Nature biotechnology 30: 521–530.

Singh A, Zhong Y, Nahlawi L, Park CS, De T, Alarcon C, Perera MA. 2020. Incorporation of DNA methylation into eQTL mapping in African Americans. In BIOCOMPUTING 2021: Proceedings of the Pacific Symposium, pp. 244–255. World Scientific.

Siraj L, Castro RI, Dewey H, Kales S, Nguyen TTL, Kanai M, Berenzy D, Mouri K, Wang QS, McCaw ZR. 2023. Functional dissection of complex and molecular trait variants at single nucleotide resolution. bioRxiv.

Sollis E, Mosaku A, Abid A, Buniello A, Cerezo M, Gil L, Groza T, Güneş O, Hall P, Hayhurst J. 2023. The NHGRI-EBI GWAS Catalog: knowledgebase and deposition resource. Nucleic acids research 51: D977–D985.

Storey JD, Bass A, Dabney A, Robinson D. 2024. qvalue: Q-value estimation for false discovery rate control. Vol R package version 2.36.0.

Tewhey R, Kotliar D, Park DS, Liu B, Winnicki S, Reilly SK, Andersen KG, Mikkelsen TS, Lander ES, Schaffner SF. 2016. Direct identification of hundreds of expression-modulating variants using a multiplexed reporter assay. Cell 165: 1519–1529.

The 1000 Genomes Project Consortium. 2015. A global reference for human genetic variation. Nature 526: 68.

Ulirsch JC, Nandakumar SK, Wang L, Giani FC, Zhang X, Rogov P, Melnikov A, McDonel P, Do R, Mikkelsen TS. 2016. Systematic functional dissection of common genetic variation affecting red blood cell traits. Cell 165: 1530–1545.

Umans BD, Battle A, Gilad Y. 2021. Where are the disease-associated eQTLs? Trends in Genetics 37: 109–124.

Urbut SM, Wang G, Carbonetto P, Stephens M. 2019. Flexible statistical methods for estimating and testing effects in genomic studies with multiple conditions. Nature Genetics 51: 187–195.

Van der Auwera GA, Carneiro MO, Hartl C, Poplin R, Del Angel G, Levy-Moonshine A, Jordan T, Shakir K, Roazen D, Thibault J. 2013. From FastQ data to high-confidence variant calls: the genome analysis toolkit best practices pipeline. Current protocols in bioinformatics 43: 11.10.11–11.10. 33.

Vockley CM, Guo C, Majoros WH, Nodzenski M, Scholtens DM, Hayes MG, Lowe WL, Reddy TE. 2015. Massively parallel quantification of the regulatory effects of noncoding genetic variation in a human cohort. Genome research 25: 1206–1214.

Võsa U, Claringbould A, Westra H-J, Bonder MJ, Deelen P, Zeng B, Kirsten H, Saha A, Kreuzhuber R, Yazar S. 2021. Large-scale cis-and trans-eQTL analyses identify thousands of genetic loci and polygenic scores that regulate blood gene expression. Nature genetics 53: 1300–1310.

Wang Y-Y, Lin S-Y, Liu P-H, Cheung BM, Lai W-A. 2004. Association between hematological parameters and metabolic syndrome components in a Chinese population. Journal of Diabetes and its Complications 18: 322–327.

Waterland RA, Jirtle RL. 2003. Transposable elements: targets for early nutritional effects on epigenetic gene regulation. Molecular and Cellular Biology 23: 5293–5300.

Weaver IC, Cervoni N, Champagne FA, D’Alessio AC, Sharma S, Seckl JR, Dymov S, Szyf M, Meaney MJ. 2004. Epigenetic programming by maternal behavior. Nature neuroscience 7: 847–854.

Yim YY, Teague CD, Nestler EJ. 2020. In vivo locus-specific editing of the neuroepigenome. Nature Reviews Neuroscience 21: 471–484.

Yin Y, Morgunova E, Jolma A, Kaasinen E, Sahu B, Khund-Sayeed S, Das PK, Kivioja T, Dave K, Zhong F. 2017. Impact of cytosine methylation on DNA binding specificities of human transcription factors. Science 356: eaaj2239.

Zeng B, Bendl J, Kosoy R, Fullard JF, Hoffman GE, Roussos P. 2022. Multi-ancestry eQTL meta-analysis of human brain identifies candidate causal variants for brain-related traits. Nature genetics 54: 161–169.

Zeng Y, Jain R, Lam M, Ahmed M, Guo H, Xu W, Zhong Y, Wei G-H, Xu W, He HH. 2023. DNA methylation modulated genetic variant effect on gene transcriptional regulation. Genome Biology 24: 285.

Zou Z, Ohta T, Oki S. 2024. ChIP-Atlas 3.0: a data-mining suite to explore chromosome architecture together with large-scale regulome data. Nucleic Acids Research: gkae358.

Zuo Z, Roy B, Chang YK, Granas D, Stormo GD. 2017. Measuring quantitative effects of methylation on transcription factor–DNA binding affinity. Science advances 3: eaao1799.

